# A simple and versatile plasma membrane staining method for visualizing living cell morphology in reproductive tissues across diverse plant species

**DOI:** 10.1101/2025.06.11.659190

**Authors:** Yuga Hanaki, Hidemasa Suzuki, Sohta Nakamura, Sakumi Nakagawa, Keigo Tada, Hikari Matsumoto, Yusuke Kimata, Yoshikatsu Sato, Minako Ueda

**Author notes:** **Corresponding author:** Minako Ueda, **Email:**. **Author Contributions:** M.U. designed the research; Y.H., H.S., S.Nakamura, S.Nakagawa, K.T., H.M., Y.K., and Y.S. carried out the experiments; Y.H., and S.Nakamura analyzed data; and M.U. wrote the manuscript.

## Abstract

Plant reproduction involves dynamic spatiotemporal changes that occur deep within maternal tissues. In ovules of *Arabidopsis thaliana* (*A. thaliana*), one of the two synergid cells degenerates at fertilization, while the fertilized egg cell (zygote) undergoes directional elongation followed by asymmetric division to initiate embryonic patterning. However, morphological analysis of these events has been hampered by the limitations of conventional cell wall staining, which fails to label cells lacking complete walls, and by the requirement for transgenic fluorescent reporters to visualize cell outlines. Here, we report that the membrane-specific fluorescent dye FM4-64 readily permeates ovules, allowing clear visualization of reproductive cell morphology both before and after fertilization. This staining method supports high-resolution time-lapse imaging and quantitative analysis of early embryogenesis in living tissues. Importantly, it is applicable not only to the angiosperm *A. thaliana* but also to the liverwort *Marchantia polymorpha* (*M. polymorpha*) and the fern *Ceratopteris richardii* (*C. richardii*), enabling the visualization of live reproductive cell structures within maternal tissues and revealing fertilization-associated morphological changes. This simple and robust method thus provides a valuable tool for spatiotemporal and quantitative analyses of reproductive processes across a broad range of plant species, without the need to generate transgenic lines.

## Introduction

Sexual reproduction is a fundamental biological process that ensures the continuity, diversity, and evolution of plants. In most land plants, egg cells (female gametes) develop deep within maternal tissues, where fertilization occurs following the arrival of sperm cells (male gametes), triggering the onset of embryogenesis. Traditionally, plant reproductive cells have been studied using fixed samples stained to visualize cell walls (Jensen, 1965, Mansfield and Briarty, 1991, Johnson and Renzaglia, 2008). However, because egg cells fuse with sperm cells at the plasma membrane (PM), they often lack complete cell walls and are therefore particularly sensitive to fixation-induced stress and are poorly visualized with conventional cell wall stains (Musielak *et al*., 2015, Tofanelli *et al*., 2019, Hisanaga *et al*., 2021). Consequently, high-resolution imaging of intact, living reproductive cells typically requires the generation of transgenic lines expressing fluorescent markers, limiting such studies to a small number of model species.

In the widely used angiosperm model *Arabidopsis thaliana* (*A. thaliana*), the egg cell resides adjacent to two synergid cells within the ovule. Upon recognition of the pollen tube, one synergid cell degenerates to release sperm cells into the ovule, and the remaining synergid cell is eliminated after fertilization (Maruyama *et al*., 2015). Whole-mount imaging techniques that combine tissue clearing methods with fluorescent probes and confocal microscopy have been developed to visualize ovule structure in three dimensions while preserving endogenous fluorescent proteins (Musielak *et al*., 2015, Park *et al*., 2019, Tofanelli *et al*., 2019, Kurihara *et al*., 2021, Attuluri *et al*., 2022). For example, the ClearSee clearing method, in combination with cell wall-specific dyes such as SCRI Renaissance 2200 (SR2200) and Calcofluor White (CFW), enables detailed imaging of developing ovules that express fluorescent reporters (Tofanelli *et al*., 2019). However, these techniques are unsuitable for visualizing egg and synergid cells, which lack complete cell walls (Musielak *et al*., 2015, Tofanelli *et al*., 2019, Attuluri *et al*., 2022).

Comparable limitations exist in non-angiosperm species. In the liverwort *Marchantia polymorpha* (*M. polymorpha*), SR2200 staining highlights maternal tissues but fails to label egg cells or young zygotes within the archegonia (Hisanaga *et al*., 2021). In ferns such as *Ceratopteris richardii* (*C. richardii*) and its congener *C. thalictroides*, histological sectioning has revealed three-dimensional archegonial organization and patterns of zygotic division (Johnson and Renzaglia, 2008, Lopez-Smith and Renzaglia, 2008, Cao *et al*., 2009, Cao *et al*., 2010a, Cao *et al*., 2010b). However, the morphology of living reproductive cells remains unexplored.

In parallel, *in vitro* ovule cultivation systems have been established in *A. thaliana*, enabling live-cell imaging of reproductive structures (Gooh *et al*., 2015, Kurihara *et al*., 2017, Ueda *et al*., 2020). Combining fluorescent reporters for intracellular structures with two-photon excitation microscopy (2PEM) has allowed researchers to visualize dynamic zygotic processes such as vacuole migration along actin filaments (Kimata *et al*., 2016, Kimata *et al*., 2019). PM-specific transgenic reporters have enabled visualization of egg cells, synergid cells, and early zygotes (Susaki *et al*., 2021, Kang *et al*., 2023). Furthermore, image analysis techniques have permitted quantitative assessment of growth and subcellular dynamics (Hiromoto *et al*., 2023, Kang *et al*., 2023, Kang *et al*., 2024b). Nonetheless, live imaging remains labor-intensive and limited to transgenic model plants.

To facilitate non-transgenic live-cell visualization, membrane-specific fluorescent dyes such as propidium iodide (PI) and FM4-64 (N-(3-triethylammoniumpropyl)-4-(p-diethylaminophenyl-hexatrienyl) pyridinium dibromide) have been used to label living plant cell membranes (Bolte *et al*., 2004, Otero *et al*., 2016, Canher *et al*., 2022). While these dyes have been used to label developing ovules and isolated embryos (Rademacher *et al*., 2011, Figueiredo *et al*., 2016), their utility for labeling living reproductive cells within intact ovules remains poorly characterized.

In this study, we demonstrate that a combination of *in vitro* ovule cultivation, FM4-64 staining, and 2PEM enables clear visualization of living egg cells, zygotes, and early embryos in *A. thaliana*, preserving cellular architecture within ovules. We further show that this approach is applicable to *M. polymorpha* and *C. richardii*, facilitating simultaneous imaging of reproductive cells and surrounding maternal tissues. This versatile and accessible method provides a valuable platform for quantitative and spatiotemporal analysis of reproductive processes across a broad phylogenetic range of plant species.

## Results

### FM4-64 dye enables visualization of living zygotes in unfixed *A. thaliana* ovules

To identify a chemical staining method suitable for visualizing living reproductive cells through ovules, we focused on zygotes of *A. thaliana* (Figure 1a). We tested three membrane dyes; FM4-64, PI, and Pontamine Fast Scarlet 4B (S4B; also known as Direct Red 23), which are known to label living tissues such as isolated embryos, root hairs, and pollen tubes (Lin and Schiefelbein, 2001, Schlereth *et al*., 2010, Park *et al*., 2011, Nagae *et al*., 2022). These dyes were applied using an established *in vitro* ovule cultivation system (Figure 1b) (Kurihara *et al*., 2017).

**Fig. 1.**
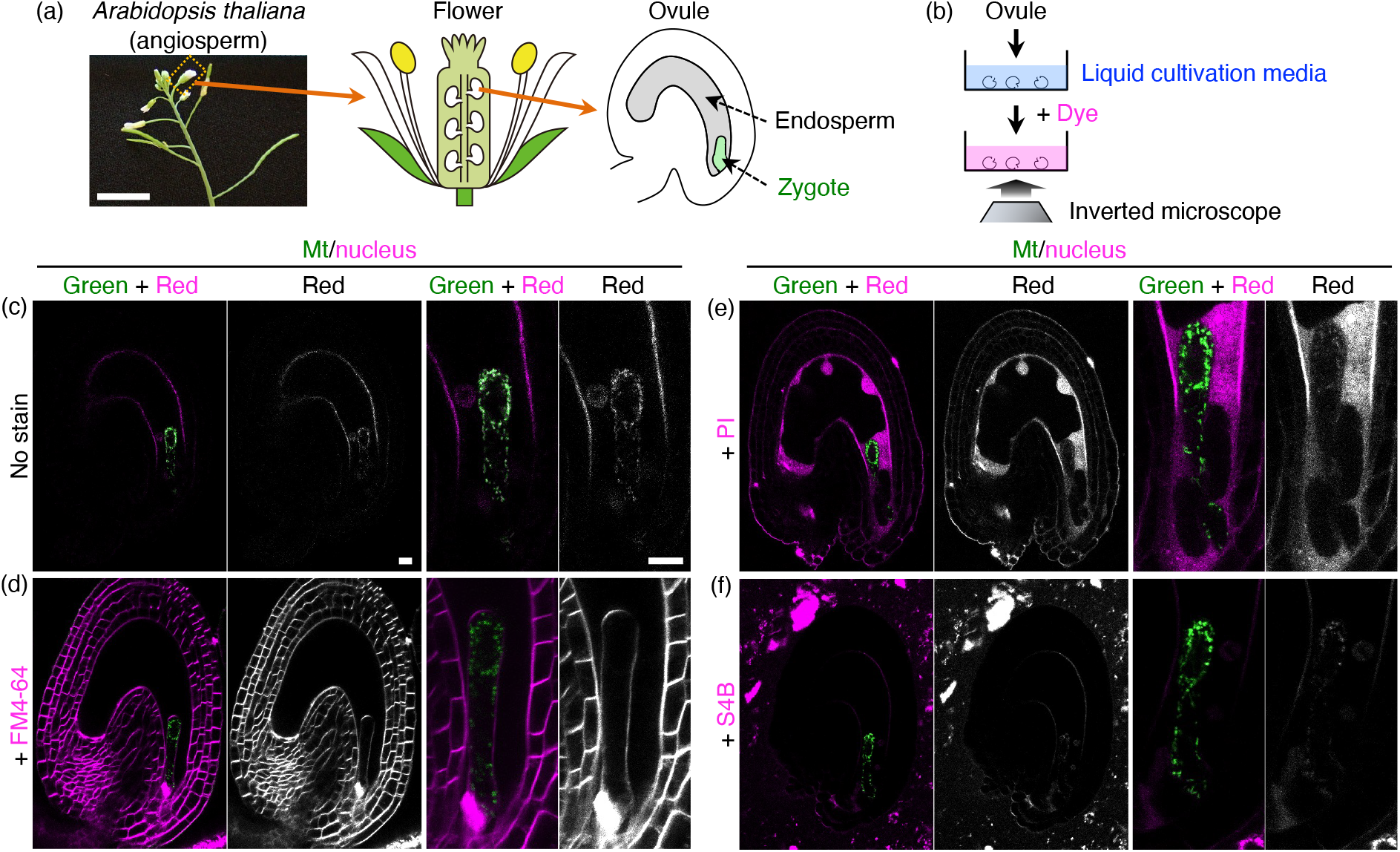
Chemical staining of zygote-containing ovules in *A. thaliana* under non-fixed conditions. (a, b) Schematic representations showing the zygote position in *A. thaliana* (a) and the chemical staining procedure using *in vitro* ovule cultivation (b). (c) Two-photon excitation microscopy (2PEM) images of zygote-containing ovules expressing the mitochondrial/nuclear (Mt/nucleus) marker. (d-f) 2PEM images of the zygotes expressing Mt/nucleus marker stained with 10 μM FM4-64 (d), 0.1 mg/L PI (e), or 0.1% S4B (f). Left panels show the merged images of the whole ovules. Right panels display green fluorescence, red fluorescence, and merged images of zygotes. Scale bars: 1 cm (a), and 10 μm (c).

To detect zygotes, we employed a dual-color mitochondrial/nuclear (mt/nuclear) marker, labelling zygotic mitochondria via DD45p::mt-Kaede and the nuclei in the zygote and endosperm via EC1p::H2B-tdTomato and DD22p::H2B-mCherry (Figure 1c-f). As previously reported, mitochondrial signals were distributed throughout the zygote, while nuclear signals localized to the apical region (Yamaoka *et al*., 2011, Kimata *et al*., 2020), and the nuclear signal was slightly masked by the mitochondrial fluorescence detected in the red channel due to the spontaneous photoconversion of Kaede during observation. The cell outlines of the ovule and zygote exhibited only faint background signals (Figure 1c). In contrast, FM4-64 staining produced a distinct outline of both the zygote and surrounding maternal cells, with signal intensity exceeding that of the photoconverted Kaede (Figure 1d). PI exhibited weak signals in the outer layers of the ovule and inside the endosperm but did not show a distinct signal outlining the zygote (Figure 1e), and S4B failed to yield detectable fluorescence (Figure 1f). These results show that FM4-64 effectively permeates living ovules and highlights the boundaries of both zygotic and maternal tissues.

### FM4-64 staining effectively visualizes living egg cells, synergid cells, and early embryos

Next, we examined whether FM4-64 staining enables the visualization of living reproductive cells and embryos at various developmental stages through ovules (Figure 2a). For comparison, we first observed fixed unfertilized ovule, which was cleared using ClearSeeAlpha and stained with the cellulose-specific dye CFW (Figure 2b). The microtubule (MT) and nucleus of the egg cell were specifically labeled with EC1p::Clover-TUA6 and ABI4p::H2B-tdTomato (MT/nucleus), respectively. This fixed sample exhibited distorted egg cell shape and diffuse MT signals, without clear fiber structures. CFW staining was restricted to the basal (micropylar) region of the egg cell, and also at the filiform apparatus of the synergid cells, where plasma membranes and cell walls are densely packed (Leshem *et al*., 2013). These observations confirmed that these cells lacking complete cell walls cannot be fully visualized using CFW staining.

**Fig. 2.**
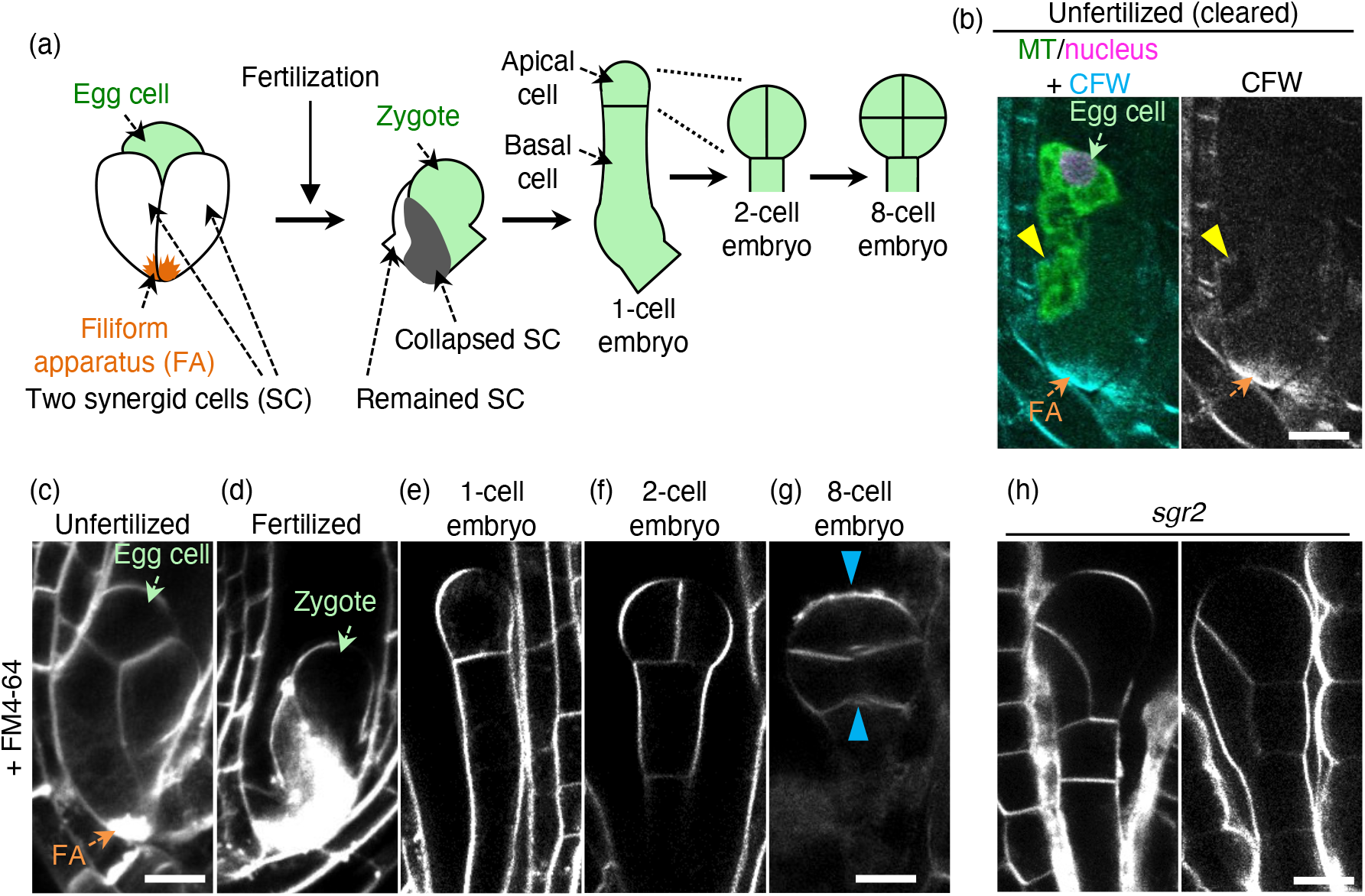
FM4-64 staining of *A. thaliana* ovules at various developmental stages. (a) Schematic illustrations of cellular arrangement before and after fertilization. (b) Cleared ovule expressing the microtubule/nuclear (MT/nucleus) marker. Cell walls were stained with CFW following PFA fixation and ClearSeeAlpha clearing. Yellow arrowheads indicate CFW signals at the basal micropylar region of the egg cell, and orange arrow shows filiform apparatus (FA). (c-g) 2PEM images of FM4-64-stained ovules from non-transgenic wild-type plant at the indicated stages. Blue arrowheads in (g) indicate unstained cell boundaries. (h) 2PEM images of FM4-64-stained ovules from non-transgenic *sgr2* mutants. Scale bars: 10 μm.

In contrast, FM4-64 staining clearly visualized the complete outlines of both the egg cell and synergid cells including filiform apparatus (Figure 2c and Movie S1). After fertilization, FM4-64 staining enabled clear visualization of zygotes and surrounding cells (Figure 2d and Movie S1). Notably, a dense fluorescent signal likely represented a collapsed synergid cell that had fully absorbed the FM4-64 dye.

Following asymmetric division of the elongated zygote, a 1-cell embryo was formed. The apical cell continued to divide, giving rise to 2-to 8-cell stage embryos. FM4-64 staining successfully visualized cell outlines throughout these stages (Figure 2e–g and Movie S1). However, at the 8-cell stage, some division planes showed weak or no staining, potentially due to reduced dye penetration as ovule development progressed. Despite these limitations, FM4-64 staining was sufficient for the precise observation of early embryo patterning, as we detected abnormal cell division planes in *shoot gravitropism2* (*sgr2*) mutant, which exhibits impaired vacuolar dynamics and abnormal embryonic patterning (Figure 2h) (Kato *et al*., 2002, Kimata *et al*., 2019).

These observations demonstrate that FM4-64 staining provides reliable and accurate visualization of the morphology of egg cells, synergid cells, zygotes, and early embryos—even in the absence of complete cell walls.

### FM4-64 staining is sufficient for quantitative live-cell imaging

We next investigated whether FM4-64 staining can be applicable to time-lapse observation (Figure 3). To quantitatively compare the growth dynamics of stained and unstained ovules, we focused on the apical cell at the 1-cell embryo stage. We recently reported that the apical cell nucleus is retained at the cell bottom, followed by the emergence of a new cell plate from the same side, leading to vertical division (Tanaka *et al*., 2024). This cell division dynamics was confirmed using a dual-color PM/nuclear marker, consisting of a PM reporter (EC1p::Clover-SYP121) and the above-mentioned nuclear reporters (EC1p::H2B-tdTomato and DD22p::H2B-mCherry) (Figure 3a and Movie S2).

**Fig. 3.**
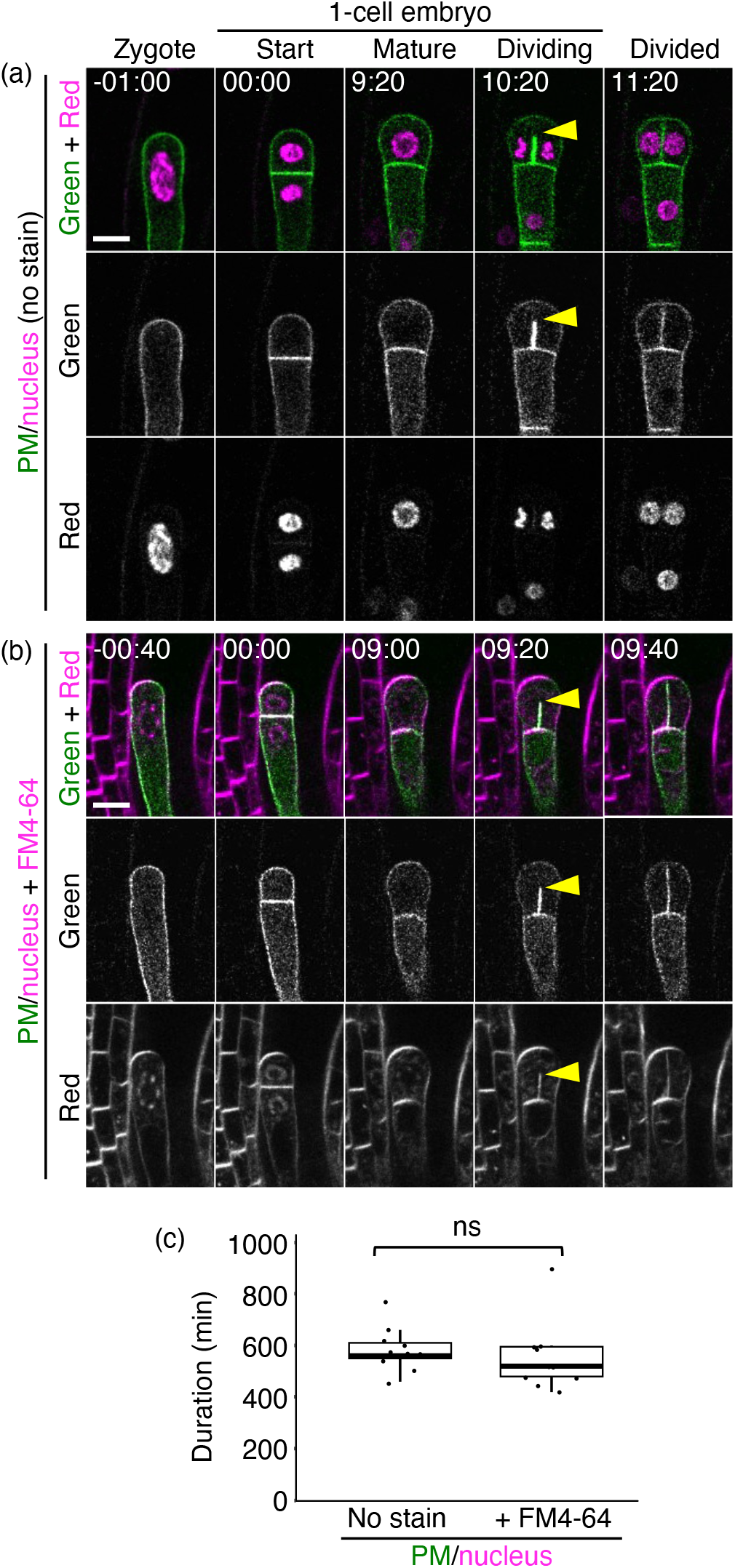
Live-cell imaging of FM4-64-stained *A. thaliana* ovules. (a and b) Time-lapse 2PEM images of wild-type ovules expressing a PM/nuclear marker in the absence (a) or presence (b) of 10 μM FM4-64. Numbers indicate the time (h:min) from zygote division. Green and red fluorescence images, along with the merged images, are presented. Arrowheads indicate elongating edges of forming cell plates in dividing apical cells. (c) Duration of apical cell division with or without 10 μM FM4-64. No significant difference (ns) was detected by the Wilcoxon rank-sum test (p = 0.26; n = 11 and 10 for control and 10 μM FM4-64-stained ovules, respectively). Scale bars: 10 μm.

A similar apical cell behavior was observed in FM4-64-stained ovules, although the red nuclear signal was slightly masked by the bright FM4-64 fluorescence (Figure 3b and Movie S3). Small, diffuse cytoplasmic dots were also observed, likely representing the vesicles internalizing FM4-64. Importantly, FM4-64 signal co-localized with the PM reporter at emerging cell plates, confirming that FM4-64 effectively labels nascent membranes without temporal delay, consistent with findings in tobacco BY-2 cells (Bolte *et al*., 2004). Furthermore, the duration from apical cell formation (zygotic division) to vertical division was indistinguishable in control and FM4-64-stained ovules (Figure 3c).

These results confirm that FM4-64 staining does not impair early embryo development and is well suited for high-resolution, quantitative live-cell imaging.

### FM4-64 staining enables visualization of reproductive cells of *M. polymorpha*

We next tested whether FM4-64 staining is applicable to non-angiosperms by examining its utility in the liverwort *M. polymorpha*, which diverged from angiosperms over 510 million years ago (Figure 4a) (Morris *et al*., 2018). In *M. polymorpha*, eggs and zygotes form within archegonia and lack rigid cell walls (Figure 4b) (Hisanaga *et al*., 2021, Bao *et al*., 2024). To assess the structural fragility in fixation, we observed cleared archegonia using the ClearSeeAlpha method, combined with an egg-and zygote-specific nuclear envelope marker (MpECpro:MpSUN-GFP) and nuclear staining with the DNA dye Kakshine PC1 (Kakshine®Yellow) (Hisanaga *et al*., 2021, Kurihara *et al*., 2021, Uno *et al*., 2021). As previously reported, these samples showed distorted nuclei and flattened cellular morphology (Figure 4c) (Hisanaga *et al*., 2021). One day after fertilization (1 DAF), zygotes contained both female pronucleus and a smaller male pronucleus, and these, along with the overall cell shape, also showed distortion (Figure 4c).

**Fig. 4.**
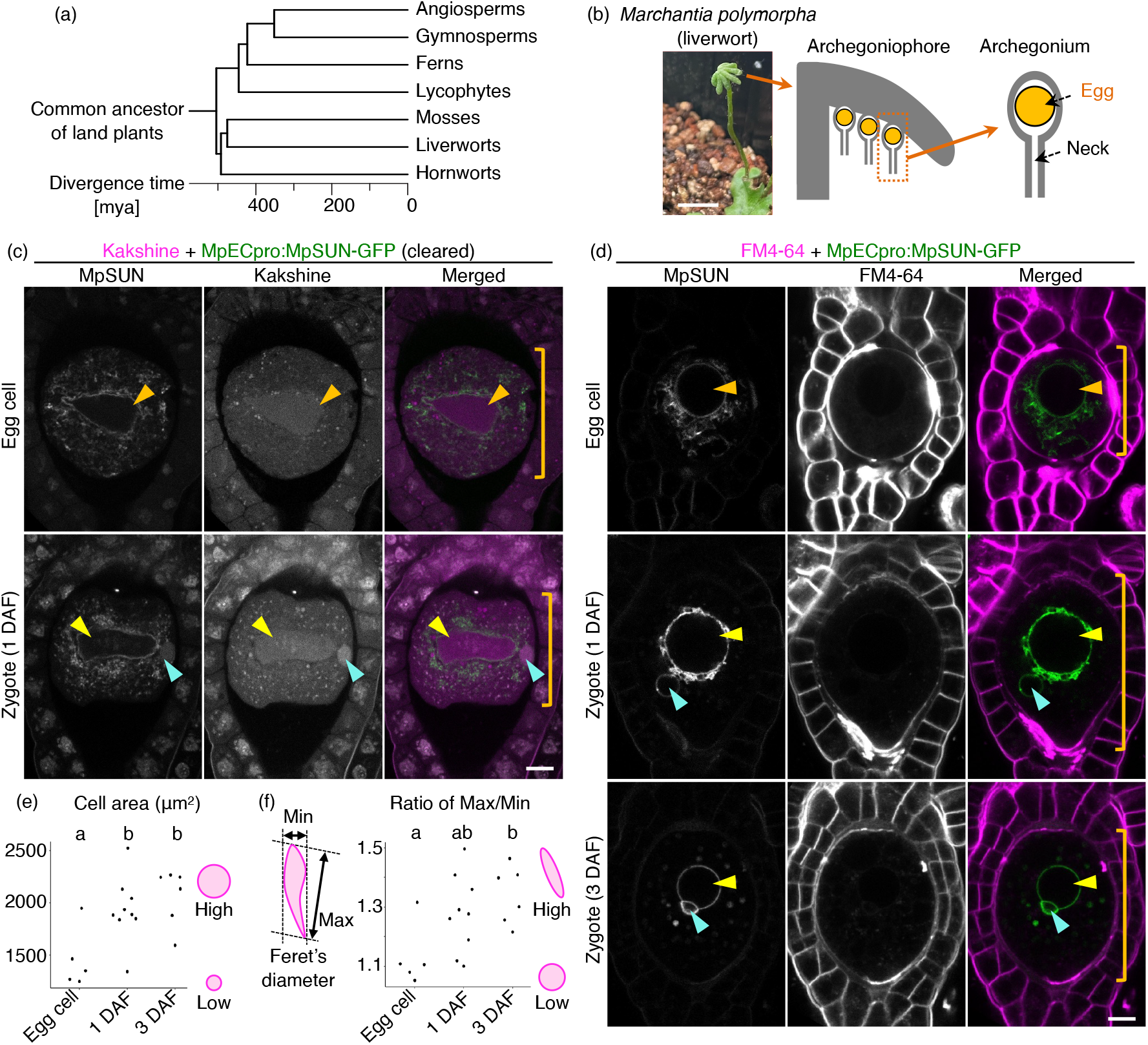
Visualization of eggs and zygotes in *M. polymorpha*. (a) Phylogenetic tree of land plants. (b) Schematic representation showing the egg position in *M. polymorpha*. (c) 2PEM images of cleared archegonia expressing the MpECpro:MpSUN-GFP marker at the indicated stages. Nuclei were stained with Kakshine following PFA fixation and ClearSeeAlpha clearing. Orange, yellow, and cyan arrowheads indicate the egg nucleus, female pronucleus, and male pronucleus, respectively. Orange brackets indicate the cell size of the egg and zygote. (d) 2PEM images of FM4-64-stained archegonia expressing the MpECpro:MpSUN-GFP marker at the indicated stages. (e, f) Quantification of cell area (e) and the ratio of Max/Min of Feret’s diameter of cell (f). The left illustration in (f) schematically defines Feret’s diameter, and the right illustrations of the graphs show the correlations between cell shape features and the corresponding values. Different lowercase letters indicate statistically significant differences (Tukey-Kramer test; p < 0.05; n = 5, 8 and 6 for egg, and zygotes at 1 and 3 DAF, respectively). Scale bars: 1 cm (b), and 10 μm (c and d).

By contrast, FM4-64 staining of living archegonia revealed well-preserved cellular architecture. Both eggs and zygotes exhibited spherical nuclei, except for male pronuclei at 3 DAF, which were tightly attached to the female pronucleus (Figure 4d). This spherical nuclear morphology was consistent with previous observations in living cells (Hisanaga *et al*., 2021), indicating minimal perturbation of intracellular structures.

Remarkably, FM4-64 staining enabled visualization of cell shape changes: eggs appeared spherical, while zygotes at 1 and 3 DAF adopted a teardrop-like morphology that filled the inner archegonial cavity (Figure 4d). This cell enlargement and deformation were quantified as significant increases in cell area (Figure 4e) and in the Max/Min ratio of Feret’s diameter, an index of cellular aspect ratio (Figure 4f). The discrepancy in morphology between living and fixed samples highlights the fragility of cells lacking rigid walls. Moreover, FM4-64 staining allowed concurrent visualization of surrounding maternal tissue architecture, confirming its broad utility in *M. polymorpha*.

### FM4-64 staining enables imaging of reproductive structures in *C. richardii*

To further assess the applicability of FM4-64 staining, we examined the model fern *C. richardii* (Figures 4a and 5a). In this species, multiple archegonia containing eggs form near the meristematic region of the hermaphroditic gametophyte (Figure 5b and 5c) (Johnson and Renzaglia, 2008, Lopez-Smith and Renzaglia, 2008). Sperms enter the archegonium through the region where neck canal cell (NCC) and ventral canal cell (VCC) occupy and then get into the egg. The resulting zygote undergoes a vertical division toward the meristematic region and proceeds embryo patterning (Figure 5b). These spatial patterns were observed only using fixed tissues (Johnson and Renzaglia, 2008, Lopez-Smith and Renzaglia, 2008), but living cell architecture were not known. In the Z-series of FM4-64-stained unfertilized archegonium, three-dimensional composition of NCC (Figure 5d), VCC (Figure 5e), and egg (Figure 5f–h) were clearly visualized through whole gametophyte (Movie S4). Furthermore, a small hole-like less-stained region was found on the egg surface (Figure 5f), resemmbling the fertilization pore found in other fern, *C. thalictroides*, from which sperms enter (Cao *et al*., 2010a). In addition, we observed the unstained region in egg inside (Figure 5f-i), which would be the cup-shaped nucleus found in the maturing egg cell of *C. thalictroides* (Cao *et al*., 2010a). After fertilization, FM4-64 staining visualized the zygote division direction toward the meristematic region (Figure 5j and k). The 2-and 4-cell embryos showed a bit larger size compared to the unfertilized egg, and nuclei were detected as the unstained region (Figure 5l and m, compare to Figure 5f-h). In 4-cell embryo, some cell division planes exhibited either weak or no signal (Figure 5m), consistent with observations in *A. thaliana* (Figure 2g).

**Fig. 5.**
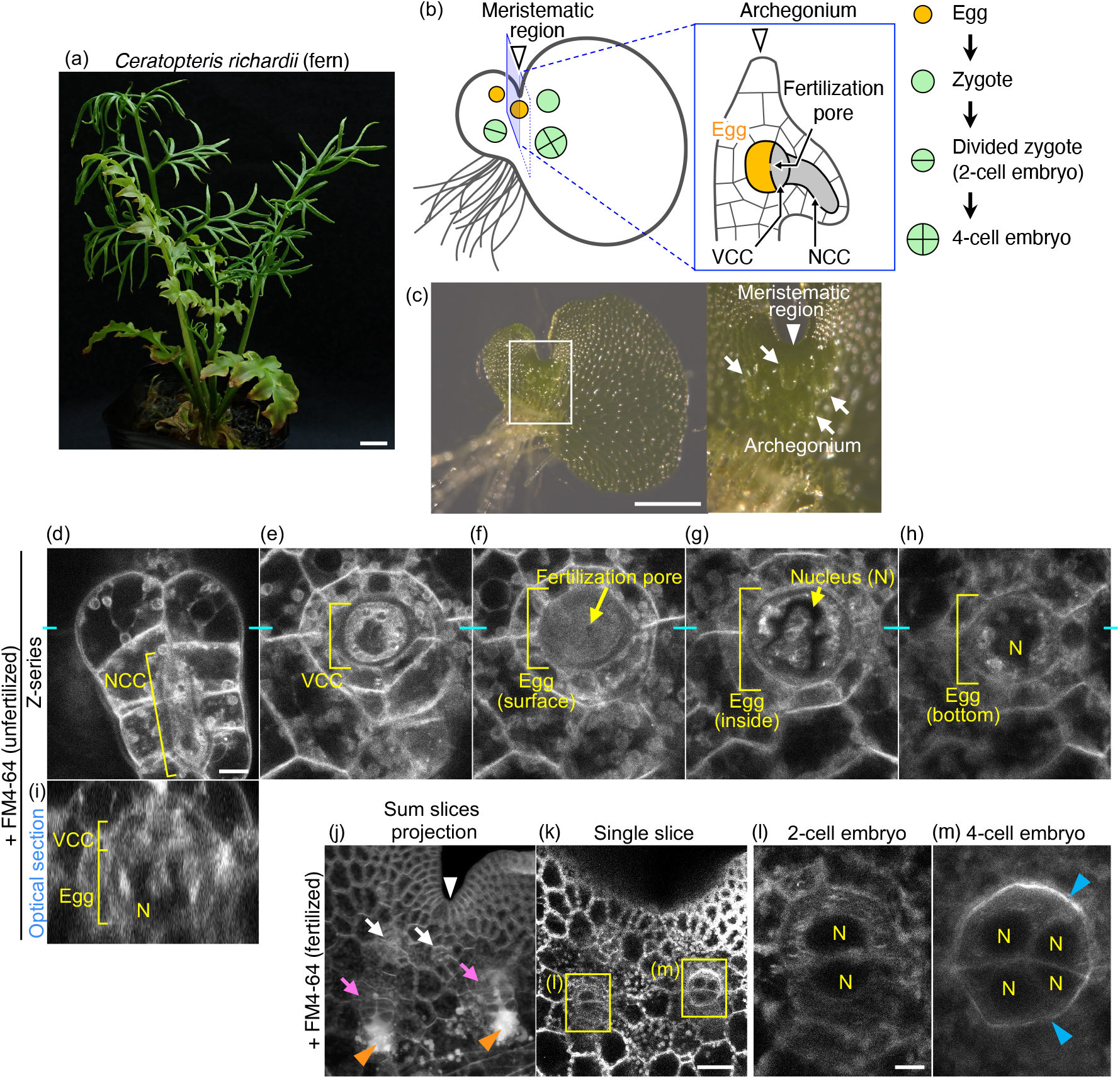
Visualization of reproductive structures and early embryos in *C. richardii*. (a) Sporophyte of *C. richardii*. (b) Schematic representation of the hermaphrodite gametophyte, archegonium, egg, zygote, and early embryo. The left illustration shows the egg, zygote, and embryos positioned around the meristematic region (white arrowhead) of the gametophyte. The illustration enclosed by a blue square depicts a cross-section of the archegonium. The right illustration shows the cell division pattern of the zygote. NCC: neck canal cell, VCC: ventral canal cell. (c) Bright-field image of hermaphrodite gametophyte. The right panel shows a magnified view of the region enclosed by a white square. The white arrowhead and white arrow indicate the meristematic region and archegonium, respectively. (d–h) Z-series of the FM4-64-stained unfertilized archegonium. Different focal planes including the NCC (d), VCC (e), and the surface (f), inner-(g) and bottom- (h) layers of the egg, respectively. N: nucleus. (i) An optical cross section image along the position indicated by cyan lines in (d–h). (j, k) FM4-64-stained meristematic region of a fertilized gametophyte. Sum slices projection reconstructed from a z-series (j) and a single slice (k) are shown. White and magenta arrows indicate the unfertilized and fertilized archegonia, respectively. White and orange arrows indicate the meristematic region and accumulated sperms, respectively. (l, m) 2- and 4-cell embryos, which are indicated by yellow squares in (k). Blue arrowheads indicate unstained cell boundaries. Scale bars: 1 cm (a), 500 μm (c), 50 μm (k), and 10 μm (d and l).

These findings demonstrate that FM4-64 staining is effective for visualizing living gametophyte structures, gametes, and early embryonic development in ferns.

## Discussion

In this study, we demonstrated that FM4-64 staining is a powerful and versatile method for high-resolution, three-dimensional visualization of reproductive tissues in living plants, enabling the simultaneous observation of both surrounding mother tissues and the internally hidden gametic cells, which lack rigid cell walls. This technique enables accurate imaging of gametes, zygotes, and early embryos without relying on transgenic fluorescent markers, which have previously limited such studies to a few model species. This ability not only accelerates the detailed characterization of newly identified mutants but also low-fertility mutants and non-model plant species where transgenic techniques are currently unavailable.

We validated the utility of FM4-64 staining in three phylogenetically distant species: the angiosperm *A. thaliana*, the liverwort *M. polymorpha*, and the fern *C. richardii*. In *M. polymorpha*, FM4-64 staining revealed that egg and zygote nuclei exhibit spherical morphology and that zygotes progressively adopt a teardrop shape following fertilization, which could not be detected in the fixed samples (Figure 4) (Hisanaga *et al*., 2021). In *C. richardii*, we succeeded in visualizing living eggs, zygotes, and intracellular structures deep within archegonia (Figure 5). The features found in *C. richardii* are largely consistent with the previous observation of fixed samples (Johnson and Renzaglia, 2008, Lopez-Smith and Renzaglia, 2008, Cao *et al*., 2009, Cao *et al*., 2010a, Cao *et al*., 2010b). This may be because, unlike the gametes of *A. thaliana* and *M. polymorpha*, which possess no or only partial cell walls, the eggs of ferns, which restrict sperm fusion at the fertilization pore, are covered by a rigid cell wall or a thick extra egg membrane, protecting gametes during fixation (Cao *et al*., 2010a).

Although FM4-64 is a well-known membrane dye and has been used in the protonemata of the moss *Physcomitrium patens* and the pollen tubes of the gymnosperm *Picea meyeri* (Wang *et al*., 2005, van Gisbergen *et al*., 2020), it has not been widely applied to reproductive tissues. Our results demonstrate that FM4-64 is broadly applicable for visualizing reproductive cells across diverse plant lineages. With further development of *in vitro* cultivation systems, FM4-64-based imaging could facilitate time-lapse studies of zygote polarization and early embryo patterning in various species.

In *A. thaliana*, we showed that FM4-64 staining enables quantitative time-lapse observation (Figure 3). FM4-64 also visualized filiform apparatus structures and the degeneration of synergid cells after fertilization (Figure 2), indicating potential applications in imaging the fertilization process, including pollen tube attraction and sperm cell release (Hamamura *et al*., 2011). The ease of application— no fixation or washing steps—and sensitivity to abnormal division patterns highlight the potential for high-throughput mutant screening.

Moreover, the compatibility of red-fluorescent FM4-64 with green-fluorescent reporters was confirmed in *A. thaliana* (Figure 1) and *M. polymorpha* (Figure 4). This enables simultaneous imaging of cell morphology and specific intracellular processes, such as cell cycle progression (Simonini *et al*., 2024). This approach may be particularly powerful in non-transgenic species when used with green-fluorescent probes, such as MitoTracker Green and BCECF that specifically stain mitochondria and vacuole, respectively (Brauer *et al*., 1995, Poot and Pierce, 1999). FM4-64 staining may also be integrated into chemical screens to facilitate a broad range of perturbation experiments and help identify regulatory factors (Nambo *et al*., 2016, Kimata *et al*., 2023).

Despite these advantages, several limitations of FM4-64 staining should be noted. One major challenge is achieving uniform dye penetration. In *A. thaliana*, partial staining becomes more frequent during embryogenesis progression (Figure 3). This may be due to the gradual accumulation of hydrophobic barrier components, such as cutin, at the boundary between internal and external ovule tissues during ovule expansion (Coen *et al*., 2019). Penetration may be further restricted in species with thicker integuments, such as gymnosperms like *Cycas revoluta* (von Aderkas *et al*., 2022). Approaches such as pressure infiltration or the use of surfactants may enhance dye access, but conditions must be carefully optimized to avoid tissue damage (Qu *et al*., 2016, Prasad *et al*., 2018).

Another concern is that prolonged imaging may lead to internalization of FM4-64 through endocytosis, resulting in vesicular accumulation or vacuolar staining (Bolte *et al*., 2004), as observed in *A. thaliana* (Figure 3). Lowering dye concentration or washing out excess dye may mitigate this effect but could compromise signal intensity. Interestingly, this property may be advantageous for imaging organelles, as nuclei appeared as FM4-64-excluding regions in *C. richardii* embryos (Figure 5).

In conclusion, while further improvements are needed to optimize staining consistency and specificity, FM4-64 staining represents a simple and robust method for real-time, quantitative imaging of reproductive events in diverse plant species without relying on transformation. This technique holds significant potential for advancing our understanding of fertilization and embryogenesis across evolutionary lineages, particularly when integrated with genetic and pharmacological approaches.

## Materials and Methods

### Plant materials and growth conditions

All *Arabidopsis thaliana* lines used in this study were derived from the Columbia (Col-0) background. Plants were grown under long-day conditions (16 h light/8 h dark) at 18–22°C. The mitochondrial/nuclear marker line contained DD45p::mt-Kaede (Hamamura *et al*., 2011) as the mitochondria-localizing reporter and EC1p::H2B-tdTomato and DD22p::H2B-mCherry as the zygote and endosperm nuclear reporter, respectively (Kimata *et al*., 2019). The PM/nuclear marker line contained EC1p::Clover-SYP121 as the PM reporter and the above-mentioned nuclear reporter, as previously described (Kang *et al*., 2023). The MT/nuclear marker line contained EC1p::Clover-TUA6 and ABI4p::H2B-tdTomato, as previously described (Kang *et al*., 2024a). The *shoot gravitropism2-1* (*sgr2-1*) mutant was described previously (Kato *et al*., 2002).

For *Marchantia polymorpha L. subsp. ruderalis*, Takaragaike 1 (Tak-1) and Takaragaike 2 (Tak-2) accessions were used as the wild-type male and female lines, respectively. Plants were grown under continuous white light at 22 °C (Ishizaki *et al*., 2016). To induce sexual reproduction, thalli were grown on 1/2× Gamborg’s B5 agar medium under white light supplemented with far-red LED (TFL-20-FE735; Fuji Electric). Two to three weeks later, the thalli were transplanted into plastic cups containing vermiculite. The MpECpro:MpSUN-GFP marker was described previously (Hisanaga *et al*., 2021).

For *Ceratopteris richardii*, Hn-n accession was used as the wild type. Plants were grown at 28 °C under long-day conditions (16 h light/8 h dark). For sterile culture, spores were surface-sterilized as described previously (Conway and Di Stilio, 2020), and were then imbibed in distilled water for 7–10 days in the dark at room temperature to synchronize germination. To induce sexual reproduction, the spores were grown on 1/2× MS agar medium (pH 5.7) supplemented with 1% sucrose under sun light-mimic LED (SUNRAY LIGHT®; Nippon Medical & Chemical Instruments Co., LTD.). For fertilization, 10 to 12 days later, the agar plates were flooded with distilled water. To culture sporophyte plant, young sporophytes were transferred onto soil (Pure Soil Black; GEX Co.,Ltd.) supplemented with nutrients (HYPONeX; HYPONeX JAPAN CORP.,LTD.) 2–3 weeks after flooding.

### Dye staining procedures

For *A. thaliana* ovules, FM4-64 (BioTracker 640 Red C2; Sigma #SCT127) was dissolved in dimethyl sulfoxide (DMSO) to 10 mM and used at a final concentration of 10 μM in *in vitro* ovule cultivation medium (Gooh *et al*., 2015, Kurihara *et al*., 2017, Ueda *et al*., 2020). Ovules were stained for 1–3 hours at room temperature before observation. PI (Wako #169-26281) and S4B (Direct Red 23; Sigma #212490) were dissolved in distilled water to concentrations of 1 mg/mL and 10%, respectively, and applied at final concentrations of 0.1 mg/mL and 0.1% for ∼3 hours.

Fixed ovules were treated with paraformaldehyde (PFA) and cleared using ClearSeeAlpha (Kurihara *et al*., 2021). Cleared samples were stained overnight with 0.1% CFW (Wako #158067) dissolved in distilled water, and observed in ClearSeeAlpha solution.

For *M. polymorpha*, mature archegonia were inculcated in 0.1% DMSO containing 10 μM FM4-64 for ∼10 min and then transferred onto slide glass. For zygote observation, fertilized archegonia were collected 1 day after *in vitro* fertilization, as described previously (Hisanaga *et al*., 2021). For comparison, archegonia were fixed in PFA, cleared with ClearSeeAlpha, and stained with 1 μM Kakshine PC1 (Kakshine®Yellow; Cosmo Bio #KA-050Y) for ∼3 hours prior to imaging (Hisanaga *et al*., 2021, Kurihara *et al*., 2021, Uno *et al*., 2021).

For *C. richardii*, hermaphrodite gametophytes were collected within 1 day after the flooding and further cultured in *in vitro* ovule cultivation medium (Gooh *et al*., 2015, Kurihara *et al*., 2017, Ueda *et al*., 2020) containing 10 μM FM4-64 for overnight at 28 °C.

### Microscopy and image analysis

All two-photon excitation microscopy (2PEM) imaging was performed using Nikon A1 or AX inverted laser-scanning microscopes, equipped with Ti:sapphire femtosecond pulse laser (Mai Tai DeepSee; Spectra-Physics) or ultrafast tunable laser (InSight X3; Spectra-Physics), respectively. Images were acquired using a 40× water-immersion objective lens (CFI Apo LWD WI; Nikon) with Immersol W 2010 (Zeiss) or water as the immersion medium. Fluorescence signals were detected by the external non-descanned GaAsP PMT detectors. For the A1 system, two dichroic mirrors (DM495 and DM560) were used, along with a band-pass filter (525/50 nm) for green signals (Kaede, Clover, and GFP), and a mirror for red signals (tdTomato, mCherry, FM4-64, PI, S4B, and Kakshine). For the AX system, three dichroic mirrors (DM488, DM560 and DM685) were used, along with two band-pass filters (525/50 nm for green, and 605/70 nm for red).

*A. thaliana* ovule images were acquired at 25 z-stacks with 2.0-μm intervals using the above objective lens, and time-lapse images were taken every 20 min. *M. polymorpha* archegonia images were acquired using 31 or 51 z-stacks with 1.0-μm intervals for fixed and living samples, respectively. For the image quantification of *M. polymorpha* cells, the cell contours were manually traced on central plane images, and the cell area and Feret’s diameter were measured using Fiji. Low-and high-magnification images of *C. richardii* gametophytes were acquired at 61 or 81 z-stacks with 1.0-or 2.0-μm intervals, respectively. For visualization of the low-magnification *C. richardii* images, Z-projection (Sum Slices) images were generated using Fiji (https://fiji.sc/).

## Acknowledgments

We thank Hisa Yoshida, Tamiko Ambo, Satomi Watanabe, and Junko Kato for technical support, Tetsuya Hisanaga and Keiji Nakajima for providing MpECpro:MpSUN-GFP, Masaru Fujimoto for DD45p::mt-Kaede, Mitsuyasu Hasebe, Junko Kyozuka, and Ayano Go Fujimura for providing *C. richardii* spores, and Satoshi Naramoto, Dolf Weijers and Sjoerd Woudenberg for helpful discussion. This work was supported by the Japan Advanced Plant Science Network, the Japan Society for the Promotion of Science [a Grant-in-Aid for Research Activity Start-up (JP21K20650 to H.M. and JP21K20649 to H.S.), a Grant-in-Aid for Early-Career Scientists (JP22K15135 to H.M., JP23K14204 to Y.K., and JP24K18135 to H.S.), Grant-in-Aid for Transformative Research Areas (A) (JP25H01809 to Y.K.), a Grant-in-Aid for Scientific Research on Innovative Areas (JP16H06280 (Advanced Bioimaging Support)), a Grant-in-Aid for Scientific Research (B) (JP23H02494 to M.U.), JSPS KAKENHI (JP25KJ0540 to S.Nakagawa), and International Leading Research (JP22K21352) to M.U.], the Japan Science and Technology Agency [CREST (JPMJCR2121 to M.U.)], the Inamori Foundation (Inamori Research Grant to Y.K.), the Suntory Rising Stars Encouragement Program in Life Sciences (SunRiSE to M.U.), and the Toray Science Foundation (20-6102 to M.U.).

## Notes

### Competing Interest Statement

The authors have declared no competing interest.

## References

Attuluri, V.P.S., Sánchez López, J.F., Maier, L., Paruch, K. and Robert, H.S. (2022) Comparing the efficiency of six clearing methods in developing seeds of Arabidopsis thaliana. Plant Reproduction, 35, 279–293.

Bao, H., Sun, R., Iwano, M., Yoshitake, Y., Aki, S.S., Umeda, M., Nishihama, R., Yamaoka, S. and Kohchi, T. (2024) Conserved CKI1-mediated signaling is required for female germline specification in Marchantia polymorpha. Current Biology, 34, 1324-1332.e1326.

Bolte, S., Talbot, C., Boutte, Y., Catrice, O., Read, N.D. and Satiat-Jeunemaitre, B. (2004) FM-dyes as experimental probes for dissecting vesicle trafficking in living plant cells. Journal of microscopy, 214, 159–173.

Brauer, D., Otto, J. and Tu, S.-I. (1995) Selective Accumulation of the Fluorescent pH Indicator,BCECF, in Vacuoles of Maize Root-Hair Cells. Journal of plant physiology, 145, 57–61.

Canher, B., Lanssens, F., Zhang, A., Bisht, A., Mazumdar, S., Heyman, J., Wolf, S., Melnyk, C.W. and De Veylder, L. (2022) The regeneration factors ERF114 and ERF115 regulate auxin-mediated lateral root development in response to mechanical cues. Molecular plant, 15, 1543–1557.

Cao, J.-G., Wang, Q.-X. and Bao, W.-M. (2010a) Formation of the Fertilization Pore during Oogenesis of the Fern Ceratopteris thalictroides. Journal of integrative plant biology, 52, 518–527.

Cao, J.-G., Wang, Q.-X., Yang, N.-Y. and Bao, W.-M. (2010b) Cytological Events during Zygote Formation of the Fern Ceratopteris thalictroides. Journal of integrative plant biology, 52, 254–264.

Cao, J.-G., Yang, N.-Y. and Wang, Q.-X. (2009) Ultrastructure of the Mature Egg and Fertilization in the Fern Ceratopteris thalictroides. Journal of integrative plant biology, 51, 243–250.

Coen, O., Lu, J., Xu, W., De Vos, D., Péchoux, C., Domergue, F., Grain, D., Lepiniec, L. and Magnani, E. (2019) Deposition of a cutin apoplastic barrier separating seed maternal and zygotic tissues. BMC plant biology, 19, 304.

Conway, S.J. and Di Stilio, V.S. (2020) An ontogenetic framework for functional studies in the model fern Ceratopteris richardii. Developmental biology, 457, 20–29.

Figueiredo, D.D., Batista, R.A., Roszak, P.J., Hennig, L. and Köhler, C. (2016) Auxin production in the endosperm drives seed coat development in Arabidopsis. eLife, 5, e20542.

Gooh, K., Ueda, M., Aruga, K., Park, J., Arata, H., Higashiyama, T. and Kurihara, D. (2015) Live-cell imaging and optical manipulation of Arabidopsis early embryogenesis. Developmental cell, 34, 242–251.

Hamamura, Y., Saito, C., Awai, C., Kurihara, D., Miyawaki, A., Nakagawa, T., Kanaoka, M.M., Sasaki, N., Nakano, A., Berger, F. and Higashiyama, T. (2011) Live-cell imaging reveals the dynamics of two sperm cells during double fertilization in Arabidopsis thaliana. Current biology : CB, 21, 497–502.

Hiromoto, Y., Minamino, N., Kikuchi, S., Kimata, Y., Matsumoto, H., Nakagawa, S., Ueda, M. and Higaki, T. (2023) Comprehensive and quantitative analysis of intracellular structure polarization at the apical–basal axis in elongating Arabidopsis zygotes. Scientific reports, 13, 22879.

Hisanaga, T., Fujimoto, S., Cui, Y., Sato, K., Sano, R., Yamaoka, S., Kohchi, T., Berger, F. and Nakajima, K. (2021) Deep evolutionary origin of gamete-directed zygote activation by KNOX/BELL transcription factors in green plants. eLife, 10, e63399.

Ishizaki, K., Nishihama, R., Yamato, K.T. and Kohchi, T. (2016) Molecular Genetic Tools and Techniques for Marchantia polymorpha Research. Plant & cell physiology, 57, 262–270.

Jensen, W.A. (1965) The Ultrastructure and Composition of the Egg and Central Cell of Cotton. American journal of botany, 52, 781–797.

Johnson, G.P. and Renzaglia, K.S. (2008) Embryology of Ceratopteris richardii (Pteridaceae, tribe Ceratopterideae), with emphasis on placental development. Journal of plant research, 121, 581–592.

Kang, Z., Matsumoto, H., Nonoyama, T., Nakagawa, S., Ishimoto, Y., Tsugawa, S. and Ueda, M. (2023) Coordinate Normalization of Live-Cell Imaging Data Reveals Growth Dynamics of the Arabidopsis Zygote. Plant and Cell Physiology, 64, 1279–1288.

Kang, Z., Nakagawa, S., Matsumoto, H., Ishimoto, Y., Nonoyama, T., Hanaki, Y., Tsugawa, S. and Ueda, M. (2024a) Temporal changes in surface tension guide the accurate asymmetric division of Arabidopsis zygotes. bioRxiv, doi: 10.1101/2024.1108.1107.605794 (preprint).

Kang, Z., Nonoyama, T., Ishimoto, Y., Matsumoto, H., Nakagawa, S., Ueda, M. and Tsugawa, S. (2024b) A viscoelastic–plastic deformation model of hemisphere-like tip growth in Arabidopsis zygotes. Quantitative Plant Biology, 5, e13.

Kato, T., Morita, M.T., Fukaki, H., Yamauchi, Y., Uehara, M., Niihama, M. and Tasaka, M. (2002) SGR2, a phospholipase-like protein, and ZIG/SGR4, a SNARE, are involved in the shoot gravitropism of Arabidopsis. The Plant cell, 14, 33–46.

Kimata, Y., Higaki, T., Kawashima, T., Kurihara, D., Sato, Y., Yamada, T., Hasezawa, S., Berger, F., Higashiyama, T. and Ueda, M. (2016) Cytoskeleton dynamics control the first asymmetric cell division in Arabidopsis zygote. Proceedings of the National Academy of Sciences of the United States of America, 113, 14157–14162.

Kimata, Y., Higaki, T., Kurihara, D., Ando, N., Matsumoto, H., Higashiyama, T. and Ueda, M. (2020) Mitochondrial dynamics and segregation during the asymmetric division of Arabidopsis zygotes. Quantitative Plant Biology, 1, e3.

Kimata, Y., Kato, T., Higaki, T., Kurihara, D., Yamada, T., Segami, S., Morita, M.T., Maeshima, M., Hasezawa, S., Higashiyama, T., Tasaka, M. and Ueda, M. (2019) Polar vacuolar distribution is essential for accurate asymmetric division of Arabidopsis zygotes. Proceedings of the National Academy of Sciences of the United States of America, 116, 2338–2343.

Kimata, Y., Yamada, M., Murata, T., Kuwata, K., Sato, A., Suzuki, T., Kurihara, D., Hasebe, M., Higashiyama, T. and Ueda, M. (2023) Novel inhibitors of microtubule organization and phragmoplast formation in diverse plant species. Life Sci Alliance, 6.

Kurihara, D., Kimata, Y., Higashiyama, T. and Ueda, M. (2017) In Vitro Ovule Cultivation for Live-cell Imaging of Zygote Polarization and Embryo Patterning in Arabidopsis thaliana. Journal of visualized experiments : JoVE.

Kurihara, D., Mizuta, Y., Nagahara, S. and Higashiyama, T. (2021) ClearSeeAlpha: Advanced Optical Clearing for Whole-Plant Imaging. Plant & cell physiology.

Leshem, Y., Johnson, C. and Sundaresan, V. (2013) Pollen tube entry into the synergid cell of Arabidopsis is observed at a site distinct from the filiform apparatus. Plant Reprod, 26, 93–99.

Lin, Y. and Schiefelbein, J. (2001) Embryonic control of epidermal cell patterning in the root and hypocotyl of Arabidopsis. Development, 128, 3697–3705.

Lopez-Smith, R. and Renzaglia, K. (2008) Sperm cell architecture, insemination, and fertilization in the model fern, Ceratopteris richardii. Sexual plant reproduction, 21, 153–167.

Mansfield, S.G. and Briarty, L.G. (1991) Early embryogenesis in Arabidopsis thaliana. II. The developing embryo. Can J Bot, 69, 461–476.

Maruyama, D., Volz, R., Takeuchi, H., Mori, T., Igawa, T., Kurihara, D., Kawashima, T., Ueda, M., Ito, M., Umeda, M., Nishikawa, S., Gross-Hardt, R. and Higashiyama, T. (2015) Rapid Elimination of the Persistent Synergid through a Cell Fusion Mechanism. Cell, 161, 907–918.

Morris, J.L., Puttick, M.N., Clark, J.W., Edwards, D., Kenrick, P., Pressel, S., Wellman, C.H., Yang, Z., Schneider, H. and Donoghue, P.C.J. (2018) The timescale of early land plant evolution. Proceedings of the National Academy of Sciences of the United States of America, 115, E2274–e2283.

Musielak, T.J., Schenkel, L., Kolb, M., Henschen, A. and Bayer, M. (2015) A simple and versatile cell wall staining protocol to study plant reproduction. Plant Reprod, 28, 161–169.

Nagae, T.T., Takeuchi, H. and Higashiyama, T. (2022) Quantification of Species-Preferential Micropylar Chemoattraction in Arabidopsis by Fluorescein Diacetate Staining of Pollen Tubes. International journal of molecular sciences, 23, 2722.

Nambo, M., Kurihara, D., Yamada, T., Nishiwaki-Ohkawa, T., Kadofusa, N., Kimata, Y., Kuwata, K., Umeda, M. and Ueda, M. (2016) Combination of Synthetic Chemistry and Live-Cell Imaging Identified a Rapid Cell Division Inhibitor in Tobacco and Arabidopsis thaliana. Plant & cell physiology, 57, 2255–2268.

Otero, S., Desvoyes, B., Peiró, R. and Gutierrez, C. (2016) Histone H3 Dynamics Reveal Domains with Distinct Proliferation Potential in the Arabidopsis Root. The Plant cell, 28, 1361–1371.

Park, S., Szumlanski, A.L., Gu, F., Guo, F. and Nielsen, E. (2011) A role for CSLD3 during cell-wall synthesis in apical plasma membranes of tip-growing root-hair cells. Nature cell biology, 13, 973–980.

Park, Y.-G., Sohn, C.H., Chen, R., McCue, M., Yun, D.H., Drummond, G.T., Ku, T., Evans, N.B., Oak, H.C., Trieu, W., Choi, H., Jin, X., Lilascharoen, V., Wang, J., Truttmann, M.C., Qi, H.W., Ploegh, H.L., Golub, T.R., Chen, S.-C., Frosch, M.P., Kulik, H.J., Lim, B.K. and Chung, K. (2019) Protection of tissue physicochemical properties using polyfunctional crosslinkers. Nature biotechnology, 37, 73–83.

Poot, M. and Pierce, R.H. (1999) Detection of changes in mitochondrial function during apoptosis by simultaneous staining with multiple fluorescent dyes and correlated multiparameter flow cytometry. Cytometry, 35, 311–317.

Prasad, A., Sedlářová, M. and Pospíšil, P. (2018) Singlet oxygen imaging using fluorescent probe Singlet Oxygen Sensor Green in photosynthetic organisms. Scientific reports, 8, 13685.

Qu, H., Xing, W., Wu, F. and Wang, Y. (2016) Rapid and Inexpensive Method of Loading Fluorescent Dye into Pollen Tubes and Root Hairs. PloS one, 11, e0152320.

Rademacher, E.H., Möller, B., Lokerse, A.S., Llavata-Peris, C.I., van den Berg, W. and Weijers, D. (2011) A cellular expression map of the Arabidopsis AUXIN RESPONSE FACTOR gene family. The Plant Journal, 68, 597–606.

Schlereth, A., Möller, B., Liu, W., Kientz, M., Flipse, J., Rademacher, E.H., Schmid, M., Jürgens, G. and Weijers, D. (2010) MONOPTEROS controls embryonic root initiation by regulating a mobile transcription factor. Nature, 464, 913–916.

Simonini, S., Bencivenga, S. and Grossniklaus, U. (2024) A paternal signal induces endosperm proliferation upon fertilization in Arabidopsis. Science, 383, 646–653.

Susaki, D., Suzuki, T., Maruyama, D., Ueda, M., Higashiyama, T. and Kurihara, D. (2021) Dynamics of the cell fate specifications during female gametophyte development in Arabidopsis. PLoS biology, 19, e3001123.

Tanaka, S., Matsushita, Y., Hanaki, Y., Higaki, T., Kamamoto, N., Matsushita, K., Higashiyama, T., Fujimoto, K. and Ueda, M. (2024) HD-ZIP IV genes are essential for embryo initial cell polarization and the radial axis formation in Arabidopsis. Current Biology, 34, 4639-4649.e4634.

Tofanelli, R., Vijayan, A., Scholz, S. and Schneitz, K. (2019) Protocol for rapid clearing and staining of fixed Arabidopsis ovules for improved imaging by confocal laser scanning microscopy. In Plant methods, pp. 120.

Ueda, M., Kimata, Y. and Kurihara, D. (2020) Live-Cell Imaging of Zygotic Intracellular Structures and Early Embryo Pattern Formation in Arabidopsis thaliana. Methods in molecular biology (Clifton, N.J.), 2122, 37–47.

Uno, K., Sugimoto, N. and Sato, Y. (2021) N-aryl pyrido cyanine derivatives are nuclear and organelle DNA markers for two-photon and super-resolution imaging. Nature communications, 12, 2650.

van Gisbergen, P., Wu, S.Z., Cheng, X., Pattavina, K.A. and Bezanilla, M. (2020) In vivo analysis of formin dynamics in the moss P. patens reveals functional class diversification. Journal of cell science, 133.

von Aderkas, P., Little, S., Nepi, M., Guarnieri, M., Antony, M. and Takaso, T. (2022) Composition of Sexual Fluids in Cycas revoluta Ovules During Pollination and Fertilization. The Botanical Review, 88, 453–484.

Wang, Q., Kong, L., Hao, H., Wang, X., Lin, J., Samaj, J. and Baluska, F. (2005) Effects of brefeldin A on pollen germination and tube growth. Antagonistic effects on endocytosis and secretion. Plant physiology, 139, 1692–1703.

Yamaoka, S., Nakajima, M., Fujimoto, M. and Tsutsumi, N. (2011) MIRO1 influences the morphology and intracellular distribution of mitochondria during embryonic cell division in Arabidopsis. Plant cell reports, 30, 239–244.

