## Supplementary material for "A simple and versatile plasma membrane staining method for visualizing living cell morphology in reproductive tissues across diverse plant species": Movie legend

**Movie S1 (separate file).** A series of xy images of the FM4-64-stained *A. thaliana* ovules.
Optical xy stacks generated from 25 z-stack images acquired at 2.0-μm intervals. The wild-type ovules stained with 10 μM FM4-64 contains an unfertilized egg cell and embryos at the indicated developmental stages.
Scale bar: 10 μm.

**Movie S2 (separate file).** Live-cell imaging of the apical cell behavior in the unstained and FM4-64-stained *A. thaliana* ovules.
Time-lapse images of ovules expressing PM/nuclear marker, observed in the absence (upper panels) or the presence (lower panels) of 10 μM FM4-64. Images were acquired every 20 minutes. Numbers indicate the time (h:min) from zygote division.
Scale bars: 10 μm.

**Movie S3 (separate file).** A series of xy images of the FM4-64-stained *M. polymorpha* archegonia.
2PEM images of FM4-64-stained *M. polymorpha* archegonia expressing the MpECpro:MpSUN-GFP marker at the indicated stages. Optical xy stacks were generated from 51 z-stack images (egg cell and zygote at 1 DAF) and 31 z-stack images (zygote at 3 DAF), with 1.0-μm intervals.
Scale bar: 10 μm.

**Movie S4 (separate file).** A series of xy images of the FM4-64-stained *C. richardii* gametophyte.
2PEM images of FM4-64 stained *C. richardii* archegonium. Optical xy stacks were generated from 61 z-stack images with 1.0-μm intervals.
Scale bar: 10 μm.
